# Vessel compression biases red blood cell partitioning at bifurcations in a haematocrit-dependent manner: implications for tumour blood flow

**DOI:** 10.1101/2020.11.25.398297

**Authors:** Romain Enjalbert, David Hardman, Timm Krüger, Miguel O. Bernabeu

**Author notes:** Equally contributing senior authors (Timm Krüger, Miguel O. Bernabeu).

## Abstract

The tumour microenvironment is abnormal and associated with tumour tissue hypoxia, immunosuppression, and poor response to treatment. One important abnormality present in tumours is vessel compression. Vessel decompression has been shown to increase survival rates in animal models via enhanced and more homogeneous oxygenation. However, our knowledge of the biophysical mechanisms linking tumour decompression to improved tumour oxygenation is limited. In this study, we propose a computational model to investigate the impact of vessel compression on red blood cell (RBC) dynamics in tumour vascular networks. Our results demonstrate that vessel compression can alter RBC partitioning at bifurcations in a haematocrit-dependent and flowrate-independent manner. We identify RBC focussing due to cross-streamline migration as the mechanism responsible and characterise the spatiotemporal recovery dynamics controlling downstream partitioning. Based on this knowledge, we formulate a reduced-order model that will help future research to elucidate how these effects propagate at a whole vascular network level. These findings contribute to the mechanistic understanding of haemodilution in tumour vascular networks and oxygen homogenisation following pharmacological solid tumour decompression.

## Introduction

The tumour microenvironment (TME) is abnormal and associated with tumour tissue hypoxia [1], which is a known biomarker for poor prognosis [2]. In addition, tumour hypoxia is a source of immunosuppression [3] and constitutes a barrier to the success of recent promising immunotherapeutic approaches [3, 4]. As such, researchers have proposed normalising the TME to improve oxygenation and overcome these limitations [3]. One of the abnormalities of the TME is vessel compression [5, 6] which is a consequence of the proliferation of cells within a solid tumour and the growth of the tumour against its surroundings [7]. Fang *et al*. showed that the presence of compressed microvessels in tumor tissue is a promising prognosis predictor in non-small cell lung cancer patients [8]. Furthermore, Chauhan *et al*. demonstrated that normalising the TME through decompressing solid tumours leads to increased survival rate in animal models as well as increased tumour tissue oxygenation and homogeneity [9]. However, our knowledge of the biophysical mechanisms linking tumour decompression to increased tumour oxygenation is limited.

Oxygen binds to haemoglobin in red blood cells (RBCs) and is transported through the vasculature with the RBCs. Early investigations demonstrated that haematocrit (the volume fraction of RBCs in total blood) can vary between child branches of a vascular bifurcation as a function of haemodynamic and geometrical vessel properties (see [10] for a review). Our recent work has shown that vascular development works to avoid vessel segments with rare or transient RBC flow through them [11]. However, the abnormal TME is associated with marked heterogeneity in haematocrit [12], including reports of plasma-only channels [13]. We recently identified reduced inter-bifurcation distance as a source of haematocrit variation via its impact on RBC partitioning at bifurcations [14]. However, the impact that other tumour vascular phenotypes such as vessel compression play on this process is not known.

*In vitro* work has provided some evidence of how RBC suspensions behave in the presence of a compression in single, straight, channels. At low haematocrits (≤ 5%), the narrowing of a channel leads to a focussing of the RBCs towards the centre of the channel [15, 16]. Fujiwara *et al*. performed a similar experiment in an asymmetric geometry at higher haematocrits (≤ 20%) and similarly saw a focussing of the RBCs towards the channel centre. They observed that this is more pronounced at lower haematocrits [17]. The focussing of RBCs is identified to be due to an increased shear rate within the compressed section of the channel [15, 17]. However, whether the narrowing in RBC cross sectional distribution post compression is permanent or not is unclear, nor is it clear how the narrowing of the RBCs toward the channel centre has an impact on RBC partitioning at a downstream bifurcation.

Investigating the transport dynamics of oxygen and other blood-borne solutes in realistic tumour networks is challenging due to the limited experimental tools available. Several groups have proposed the use of mathematical modelling to bridge this gap [18–21]. In this work, we propose a computational model to study how vessel compression impacts the partitioning of RBCs at a downstream bifurcation. We report the novel finding that, below a critical haematocrit threshold, vessel compression alters the partitioning of RBCs at the downstream bifurcation due to a change in the cross-sectional distribution of the RBCs induced by the compression. Furthermore, we show that this is independent of flow rate and compression asymmetry. In addition, we report, for the first time, the mechanism and length scale for the cross-sectional distribution of RBCs to return to their pre-compression configuration. Finally, we propose a reduced-order model to calculate RBC partitioning at a bifurcation downstream of a compression in a computationally efficient manner. Future investigations can use this reduced-order model to link vessel compression to tumour tissue oxygen heterogeneity on a whole vascular network level.

Taken together, our findings suggest that: a) protection against abnormal partitioning at bifurcations due to naturally occurring morphological variations in vessel cross-section can be achieved during development by homogenising and increasing the average haematocrit in networks, b) increased perfusion (in terms of total flow rate through the network) is not sufficient to reverse the anomalous RBC partitioning due to vessel compression, and c) the link between solid stress and oxygen heterogeneity in tumours, and its reported reversal via stress alleviation, can be partially explained via anomalous RBC partitioning at bifurcations due to compressed vessels.

## Methods

### Physical model

We model blood flow as a suspension of deformable RBC particles in a continuous plasma phase. The plasma is treated as a continuous Newtonian fluid, with the non-Newtonian properties of blood arising from the presence of the deformable RBCs.

The model for the RBC membrane is hyperelastic, isotropic and homogeneous with a membrane energy *W* = *W* ^s^ + *W* ^b^ + *W* ^a^ + *W* ^v^ [22]. *W* ^s^ is based on Skalak’s model for the surface strain energy density [23], *W* ^b^ is a bending energy, *W* ^a^ and *W* ^v^ are penalties on changes in membrane surface area and volume, respectively [22]. We determine the deformability of the RBC by the capillary number,

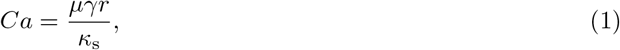

where *µ* is the fluid dynamic viscosity, *γ* is a characteristic shear rate, *r* is a characteristic length (the RBC radius) and *κ*_s_ is the strain modulus of the RBCs. The capillary number is set to 0.1, unless stated otherwise. The bending modulus is determined from a Föppl-von Kármán number of 400 for a healthy RBC [24]. The radius of the RBC is set to *r* = 4 *µ*m.

### Numerical model

We solve the fluid model with the lattice Boltzmann method (LBM) and the deformable RBC model with the finite element method (FEM). The fluid structure interaction is modelled with the immersed boundary method (IBM). This is a previously validated model from our group [22, 25]. The numerical algorithm is implemented in the software *HemeLB* (http://css.chem.ucl.ac.uk/hemelb) [26], and the simulations have been run on the ARCHER supercomputer.

Our LBM algorithm employs the D3Q19 lattice, the Bhatnagar-Gross-Krook collision operator, Guo’s forcing scheme [27], the Bouzidi-Firdaouss-Lallemand no-slip boundary condition at the walls [28], and the Ladd velocity boundary condition for inlets/outlets [29].

We discretise the RBC membrane into 720 elements and use a lattice voxel size of *0*.6667 *µ*m for the flow simulation, which is sufficient to resolve both the flow and the RBC membrane dynamics [22, 30]. We set the viscosity of the plasma and fluid inside the RBC to 1 mPa s. The density of the fluid is 1000 kg/m^3^. We chose a dimensionless lattice-Boltzmann relaxation time of 1, which gives a time step of 7.41 × 10^−8^ s.

### Geometry

We produced four geometries representing a vessel with diameter *D* and a downstream bifurcation. The first geometry is a control (no compression, Figure 1a), while the remaining three geometries feature a single compression upstream of the bifurcation. The first compression model (Figure 1b) contains a long compression without a recovery length between compression and bifurcation. In the second compression model (Figure 1c), the compression is short, and there is a short recovery length between compression and bifurcation. The last geometry (Figure 1d) features a short compression followed by a long recovery segment. The relevant geometrical parameters of all four geometries are summarised in Table 1.

**Table 1.** Dimensions of the geometry (*D* is channel diameter).

| Geometry name | compression length | distance between compression and bifurcation |
| --- | --- | --- |
| Control | N/A | N/A |
| Long compression no recovery | $4D$ | 0 |
| Short compression short recovery | $D$ | $2D$ |
| Short compression long recovery | $D$ | $25D$ |

**Figure 1.**
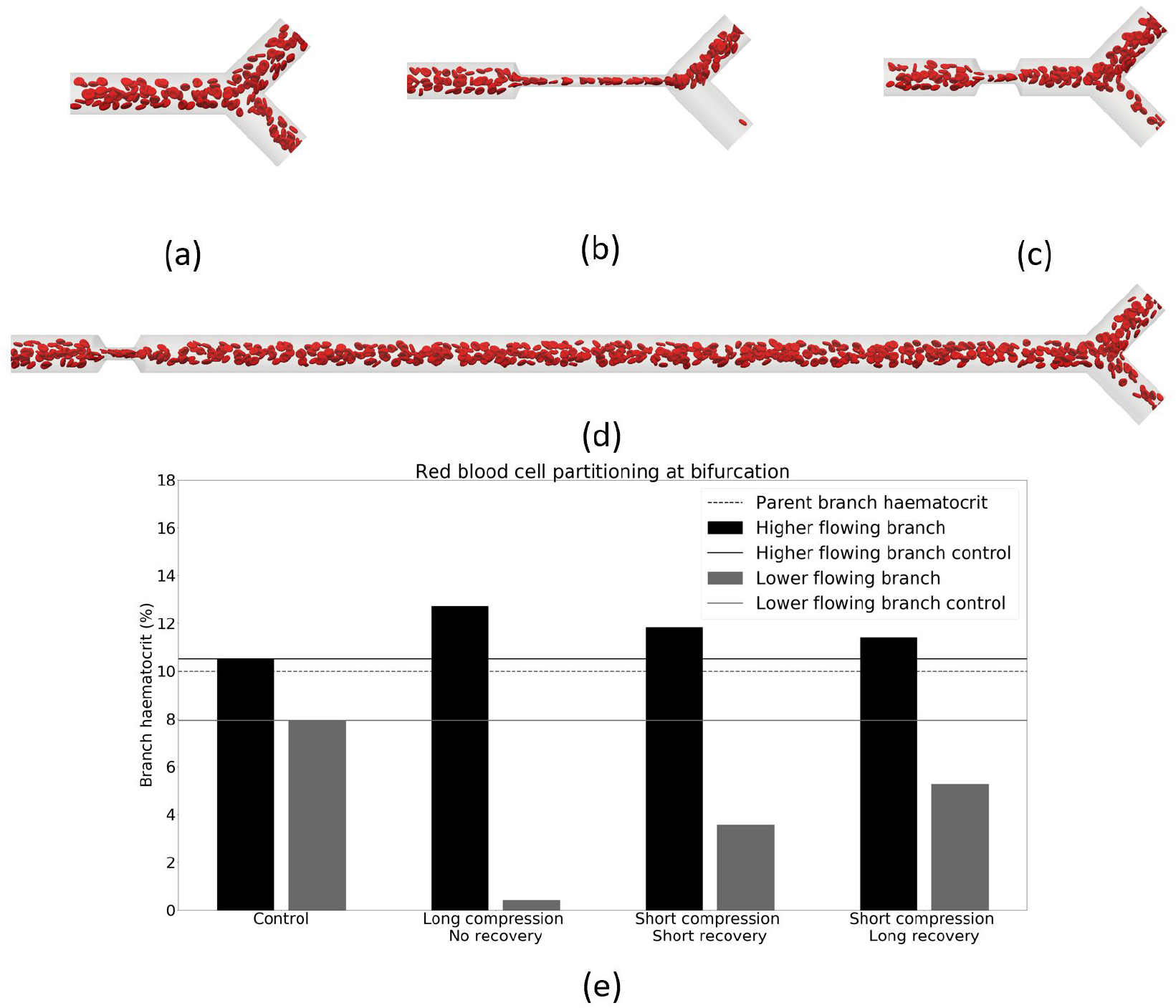
Phase separation in child branches after a bifurcation at *H*_*d*_ = 10%. (a–d) are snapshots of the control, long compression no recovery, short compression short recovery and short compression long recovery, respectively. (e) shows the haematocrit in the child branches for these four cases. Black/grey indicates the higher/lower flowing child branch, respectively. Solid lines are the control discharge haematocrits. The dotted line illustrates the discharge haematocrit of the parent branch.

We set the channel diameter to *D* = 33 *µ*m, a typical value for the tumour microvasculature [12]. We assume the cross section of the channel to be circular, except for the section that is compressed, where it takes an elliptical form. We assume the perimeter of the cross section to be constant along the channel, setting the ellipse perimeter to the same value as the uncompressed circular cross section. The segment with elliptical cross section has an aspect ratio of 4.26 [8]. The assumption of an elliptical cross-section within the compression is in line with observations from tumour histological slices where vessel compression is commonly reported as the aspect ratio of the elliptical shape of the vessel cross sections [8, 31–33].

Our aim is to focus on the effect of the compression. Therefore, we remove any effect from the slope leading to the compression by having a steep transition to and from the compression. We also remove the effect of a bifurcation asymmetry by having both child branches at the same diameter and angle from the parent branch.

### Inlet and outlet boundary conditions

We set the outflow boundary conditions at the child branches as a Poiseuille velocity profile with an imposed maximum velocity and control the ratio of these velocities such that one child branch receives 80% of the flow and the other child branch 20%. Unless specified otherwise, the inlet branch has an average velocity of 600 *µ*m/s, a typical value for the tumour microvasculature [12].

The inlet boundary condition is an arbitrary pressure value that has no impact on the simulation result. In order to reduce any memory effects and establish a quasi steady-state distribution of RBCs, the cells flow through a straight tube with a length of 25 tube diameters before entering the compression [34]; we call this length the initialisation length.

We vary the value of haematocrit within our system from 10% to 30%, covering a wide range which is physiologically present within the tumour microvasculature [12].

The physical Reynolds number of the system is 0.04. Therefore, viscous forces dominate and the system is in the Stokes flow regime. For computational tractability we set the numerical Reynolds number to 1, where inertial forces still do not play a significant role.

### Processing results

All haematocrits reported are discharge haematocrits. We calculate the discharge haematocrit, *H*_*d*_, by calculating the fraction of RBC flow to total blood flow at any channel cross section normal to the direction of flow,

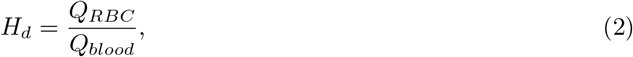

where *Q*_*RBC*_ is the volumetric flow rate of RBCs and *Q*_*blood*_ is the volumetric flow rate of blood, i.e. plasma and RBCs. The flow rate of RBCs is calculated by counting the RBCs crossing a plane normal to the direction of flow over a given period of time, Δ*t*. Knowing the volume of an RBC, *V*_*RBC*_ = 100 *µ*m^3^, one can calculate the RBC flow rate,

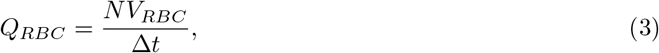

where *N* is the number of cells that have crossed the plane.

In order to quantify the distribution of the RBCs in a cross section, we measure the root mean squared distance (RMSD) of the RBC centres of mass with respect to the channel centreline,

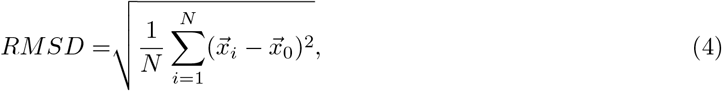

where 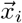 is the *i*^*th*^ RBC position, and 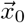 is the channel centre, both taken on a cross section normal to the direction of flow at points of interest. Figure S1 illustrates how we obtain the positions 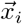 in practice.

We non-dimensionalise length by the vessel diameter *D*, unless stated otherwise. By definition, we set the downstream end of the compression as the reference point with an axial position of 0. Axial positions are positive in downstream direction (*l >* 0) and negative in upstream direction (*l <* 0).

The separatrix is an imaginary surface separating fluid particles going to one child branch from those going to the other child branch. It is an important tool for the investigation of RBC partitioning at a bifurcation [35]. We determine the separatrix by completing a simulation without RBCs to obtain streamlines unperturbed by RBCs (see Figure S2 for details).

## Results

### At 10% haematocrit, vessel compression alters RBC partitioning at a down-stream bifurcation

We start by investigating whether vessel compression has an impact on RBC partitioning at a downstream bifurcation. Simulation results for a long compression at 10% haematocrit (Figure 1b) reveal that the RBC split at the bifurcation is strongly affected by the compression. Figure 1e shows that, for a long compression, the child branch with the lower flow rate is almost depleted of RBCs and has approximately 0.5% haematocrit, whereas the control simulation indicates that the same branch has ∼ 8% haematocrit in the absence of a compression. The control simulation is in agreement with the standard plasma skimming model, see Figure S3.

The short compression short recovery geometry (Figure 1c) shows a smaller impact on the RBC split than the long compression. With ∼ 3.5% haematocrit in the child branch with the lower flow rate, this is still less than half of the haematocrit in the control simulation.

In order to test whether the symmetry of the compression has an effect on the partitioning of the RBCs at the downstream bifurcation, we investigated an asymmetric compression (Figure S4). The simulation results are similar to those obtained with a symmetric compression. Thus, we infer that, under the present conditions, the asymmetry of the compression does not have an important effect on the downstream partitioning of RBCs.

We also investigated the effect of flow rate on RBC partitioning. By changing the flow rate, we change the capillary number, which quantifies the deformation of the RBCs. We performed two additional simulations at a capillary number of 0.02 and 0.5, to cover the RBC tumbling and tank-treading regimes and the range of flow rates typical for the tumour microvasculature [12]. Figure S5 shows that the RBC partitioning does not change with capillary number, which implies that the flow rate and capillary number are not important parameters for RBC partitioning in the presence of a compression within the studied range of flow rates.

### Narrowing of cell distribution alters partitioning of RBCs

Next we investigated which mechanism leads to the observed changes in partitioning, and why the different geometries have different effects.

As blood flows through the compression, the shear rate increases since 1) the fluid velocity within the compression is larger due to mass conservation and 2) the width of the channel along the compression axis is reduced. Our simulations show that RBCs situated close to the wall prior to the compression migrate across streamlines towards the channel centre. After leaving the compression, RBCs do not immediately migrate back towards the wall. As a consequence, the RBC distribution downstream of the compression is more narrow than upstream of the compression. This explanation is in line with prior findings from experimentalists [17]. Figure 2a–d illustrates this mechanism.

**Figure 2.**
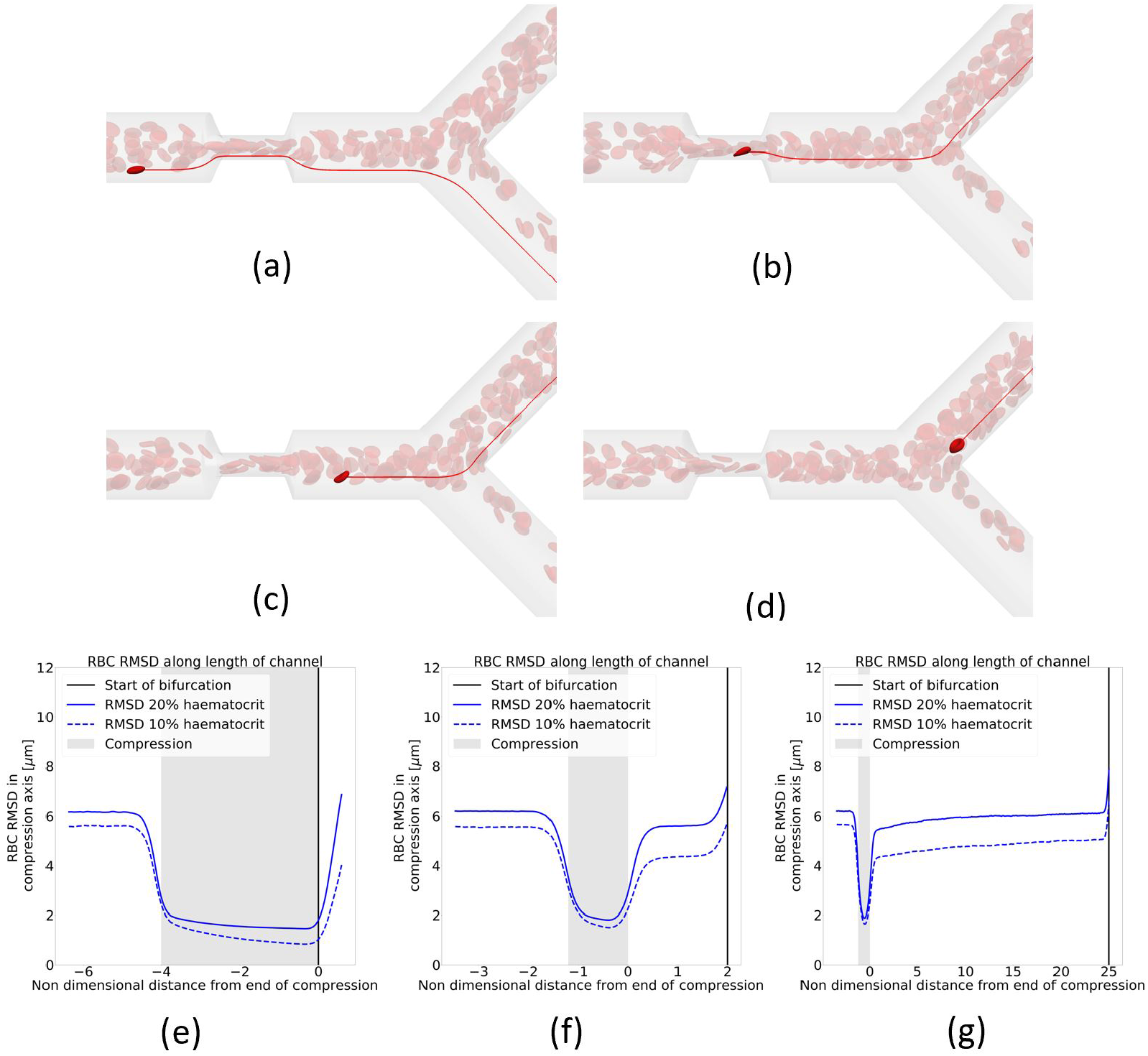
Narrowing of RBC distribution in compression. (a–d) In red are the streamlines of the underlying fluid. In bright red is an RBC of interest. (a) An RBC situated prior the compression near the vessel wall. (b) The same RBC after it has crossed streamlines within the compression. (c) The RBC exits the compression on a more central streamline than the one on which it entered the compression. (d) The RBC goes to a different branch than the pre-compression streamline it was on. (e–g) RBC RMSD along the vessel length, rigid line at *H*_*d*_ = 20% and dashed line at *H*_*d*_ = 10%. Blue line is the RMSD, black vertical line is the point of bifurcation, and shaded grey zone is the compression area. Geometries are (e) Long compression no recovery, (f) short compression short recovery, and (g) short compression long recovery. The child branch flow ratio is 4:1 in all cases.

In order to quantify the narrowing of the RBC distribution, we plot the RMSD of the RBC centres of mass along the compression axis. Figure 2e–g shows that, for all geometries, there is a narrowing of the distribution of cells in and after the compression compared to the region before the compression. However, in the short recovery geometries (Figure 2f–g), the RBC distribution partially recovers before the cells reach the bifurcation. This explains why the RBC partitioning is more affected when there is no recovery length between compression and bifurcation (Figure 2e). Previous studies report an increase of the cell free layer (CFL) post compression compared to the CFL thickness pre compression, here seen as a narrowing of the RBC distribution, which leads to a partitioning bias of RBCs towards the higher flowing branch [36].

In order to investigate the behaviour of the RBC distribution after the compression, we increased the distance between the compression and the bifurcation from 2*D* to 25*D* (Figure 1d). Figure 2g shows a gradual recovery of the RBC distribution between the compression and bifurcation, although 25 channel diameters are not sufficient to reach the same RMSD as before the compression.

While our data imply that a mechanism exists that leads to the recovery of the RBC distribution, it is not clear a priori what the underlying mechanism is. We assume that, given enough channel length after the compression, the RBC distribution will fully recover eventually. Katanov *et al*. demonstrated that, from an initially uniform distribution of RBCs in a channel, the formation of a stable CFL is governed by the shear rate time scale and takes a length of about 25 vessel diameters to form, independently of flow rate, haematocrit or vessel diameter [34]. Our data suggest that the opposite effect, the recovery of an initially heterogeneous RBC distribution where most of the RBCs are close to the channel centre, cannot be described in the same way since a length of 25 channel diameters is not sufficient for recovery. The shear rate is lower and cells move faster near the channel centreline, which should lead to a weaker shear-induced recovery of the cell distribution along a distance of 25*D*. We hypothesise that cell-cell interactions are the dominant driver for the recovery.

### Increasing haematocrit reduces bias in RBC partitioning

To test the hypothesis that cell-cell interactions drive the recovery of the distorted RBC distribution, we investigate blood flow at an increased haematocrit of 20%. Figure 3 shows that the long compression no recovery geometry still leads to a deviation from the control simulation. However, the deviation is smaller than at 10% haematocrit (Figure 1). This observation can be explained by Figure 2e–g which reveals that the narrowing of the RBC distribution is less pronounced at higher haematocrit compared to lower haematocrit. We conclude that, as the haematocrit increases, hydrodynamic cell-cell interactions become more relevant, leading to a smaller narrowing of the cell distribution by the compression as well as a faster decay of the narrowed RBC distribution after the compression.

**Figure 3.**
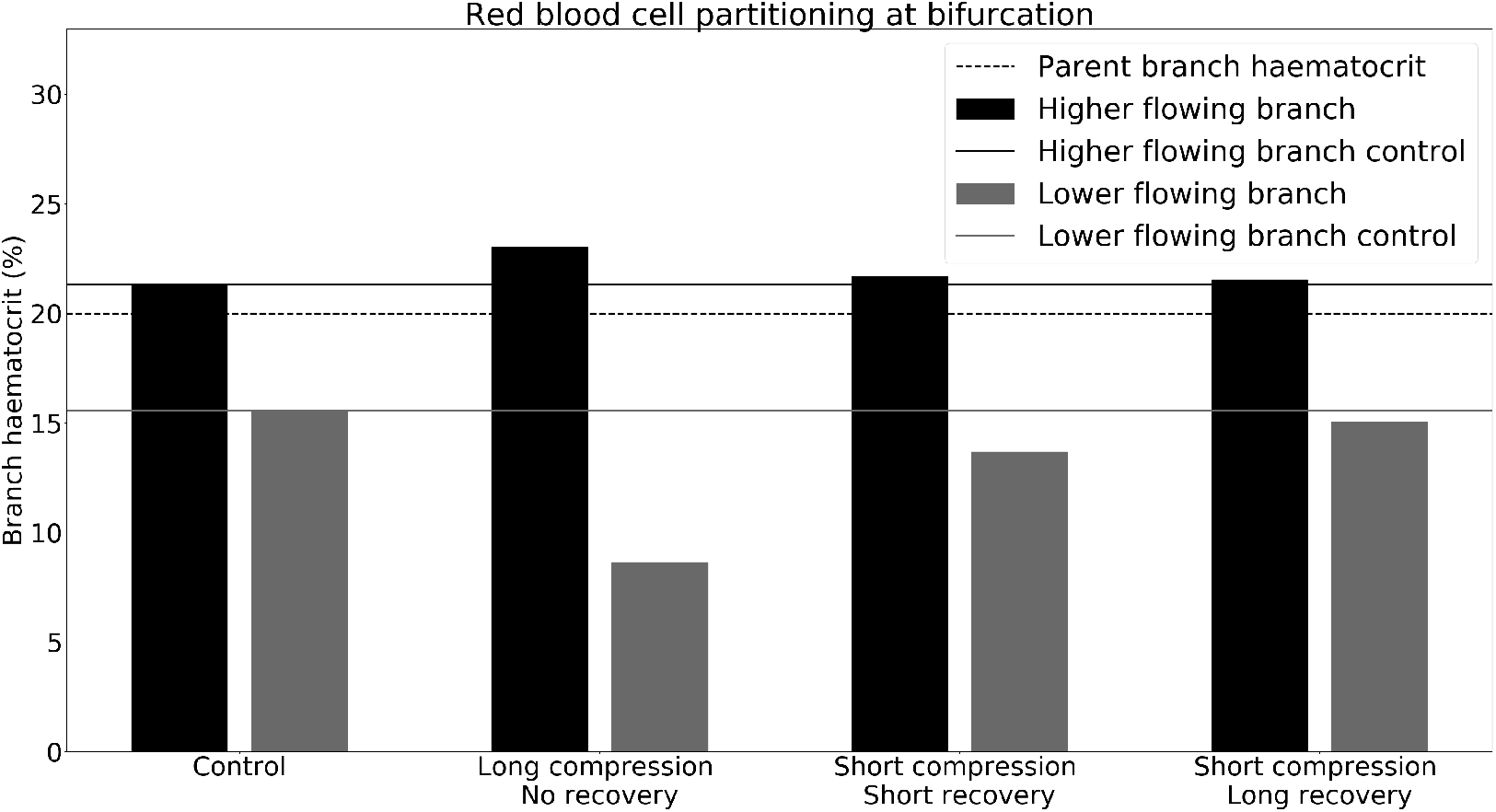
Phase separation in child branches after a bifurcation at *H*_*d*_ = 20%. Black/grey indicates the haematocrit in the higher/lower flowing child branch, respectively. Solid lines are the control discharge haematocrits. The dotted line illustrates the discharge haematocrit of the parent branch. The child branch flow ratio is 4:1 in all cases.

For the short compression long recovery geometry, the deviation from the control is almost non-existent at *H*_*d*_ = 20% (Figure 3). Whilst the control simulation shows 15.6% haematocrit in the lower flowing child branch, the compression merely alters that value to 13.7%. As can be seen from Figure 2f, not only is the narrowing of the RBC distribution in the compression smaller, but after exiting the compression the RBC distribution is much closer to its pre-compression counterpart.

We also investigated the role of the distance between the short compression and the bifurcation by increasing it from 2*D* to 25*D*. Figure 2g shows that the RBC distribution with *H*_*d*_ = 20% eventually recovers and goes back to its pre-compression level, contrary to the simulation at 10% haematocrit.

The decreasing effect of the constriction on the RBC distribution at increasing haematocrit raises the question whether there is a critical haematocrit above which the RBC partitioning is not modified by the presence of an upstream constriction. To that end, we increased the haematocrit in the parent branch to 30% and revisited the long compression geometry that has no recovery length between compression and bifurcation. Figure 4 shows that there is no significant difference in RBC split when compared to the control without compression. We conclude that the critical haematocrit value lies near 30%.

**Figure 4.**
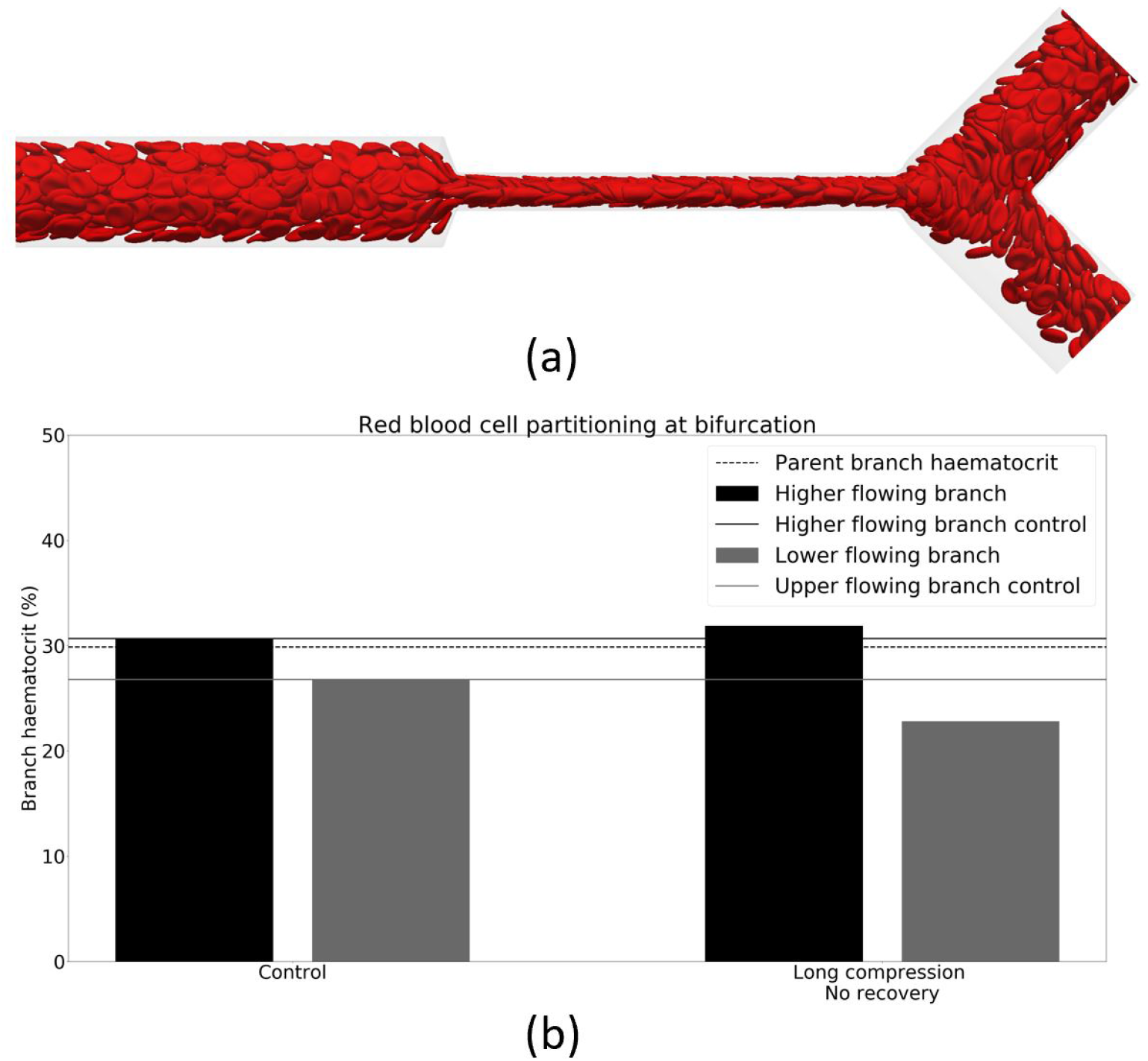
Phase separation in child branches after a long bifurcation at *H*_*d*_ = 30%. (a) shows the haematocrit of the child branches. Black/grey indicates the higher/lower flowing child branch, respectively. Solid lines are the control discharge haematocrits. The dotted line illustrates the discharge haematocrit of the parent branch. (b) shows the snapshot of the simulation in the long compression. The child branch flow ratio is 4:1.

### Reduced-order model

A challenge in the theoretical study of RBC transport in networks is computational expense [25, 37, 38]. For this reason, several authors have proposed the use of reduced-order models to quantify the partitioning of RBCs at bifurcations, which is key for tissue oxygenation modelling due to the RBC’s role as oxygen carrier. The most common model existing for partitioning of RBCs is that presented by Pries et al. [39, 40], although others exist, e.g. [41].

We have demonstrated that vessel compression indeed has an impact on the partitioning of RBCs at a downstream bifurcation and that this is not captured by a state-of-the-art reduced-order model. In order to investigate how this effect propagates on a network level, we propose a novel reduced-order model that captures this phenomenon. We make four main assumptions for the reduced-order model:

1. An RBC’s centre of mass will go to the same child branch as its underlying streamline. Similarly to reports of RBCs crossing the separatrix prior to a bifurcation [35, 36, 42], in our simulations we observed *<* 5% of RBCs near the separatrix crossing streamlines. Due to this small fraction, we deem the assumption appropriate.
2. A curved separatrix independent of the diameter ratio between the child and parent branch and independent of Reynolds number is used, whereas the curvature of the separatrix generally depends on both parameters [35, 43]. Since blood flow in the microvasculature is in the low-Reynolds regime, a small change in Reynolds number has a negligible effect on the separatrix. The diameter ratio has been shown to have a more significant impact, even at a Reynolds number of 0 [43]. However, the impact on the curvature of the separatrix is higher towards the vessel walls, where there are fewer or no RBCs due to the existence of the CFL. Considering the difficulty of parameterising a curved separatrix as a function of diameter ratios, a curved separatrix for a diameter ratio of 1 is used at the expense of a small but acceptable modelling error.
3. The cross-sectional distribution of RBC centres of mass can be approximated by a step function, whereas the distribution profile of RBCs tends to quickly, but not instantaneously, reduce at the edge of the RBC distribution as is often reported [44, 45] and indeed observed in our simulations. The step function, however, is a good fit and simplifies the model considerably. We assume the step function to take the shape of an ellipse on any given cross section of the channel (Figure 5a–c). The fraction of RBCs ending up in the top and bottom child branch is *A/*(*A* + *B*) and *B/*(*A* + *B*), respectively, defined by the areas *A* and *B* above and below the separatrix.

**Figure 5.**
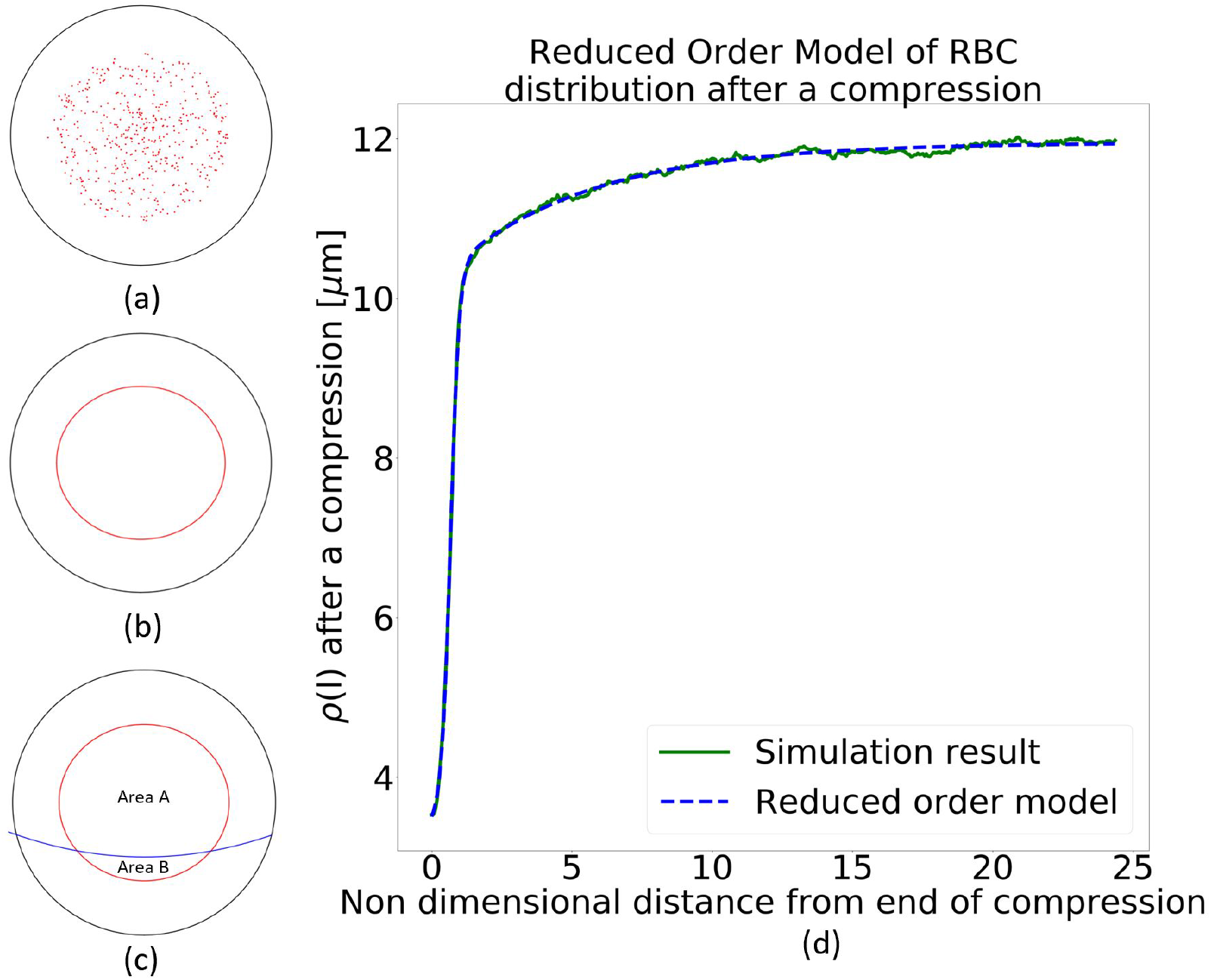
Reduced-order model. (a) The RBCs’ centres of mass are shown on a cross section. (b) An ellipse is used to represent the distribution of the RBCs. (c) The curved separatrix is added. Any RBC above the separatrix is assumed to go to the top branch, and any RBC below it to go to the bottom branch. (d) *ρ*(*l*) from the reduced-order model in Eq. (5) with parameters from Table 2 for 20% haematocrit compared to simulation data.
4. The cross section that determines which child branch an RBC enters is located about 2*D/*3 upstream of the bifurcation. Our simulations show that this is the upstream perturbation length after which the streamlines start to curve in order to enter the child branches. The length 2*D/*3 is similar to that found in other studies [35]. Therefore, the reduced-order model needs to be able to predict the cross-sectional distribution of the RBCs up to a point 2*D/*3 upstream of the bifurcation.

The step function that approximates the RBC distribution in a channel cross section has the form of an ellipse. Therefore, the major and minor semi-axes of this ellipse, *a* and *b*, need to be defined. We found that the best results are obtained when

1. the aspect ratio of the ellipse is determined by the ratio of the RMSD along the width and height directions (where the height direction is the axis of the compression),
2. the ellipse encloses 90% of the RBCs’ centres of mass.

Next, we propose a function that describes the development of the radius *ρ*, along the compression axis, of the step function along the channel length *l* between the end of the compression (defined as *l* = 0) and the point 2*D/*3 upstream of the bifurcation. The area of the ellipse is given by *A* = *πab* where *a* and *b* are the major and minor semi-axis of the ellipse for the step function. Once the minor axis *b*, which is *ρ*(*l*), and the aspect ratio *ϵ*(*l*) = *a*(*l*)*/b*(*l*) are known, the model can predict the number of RBCs entering either child branch.

Our simulations show that there are three key mechanisms governing the lateral RBC distribution when entering and leaving the compression. The first is that the RBC distribution is suddenly narrowed by the compression. Secondly, upon exiting the compression, the RBC distribution sees a quick but only partial lateral recovery due to the expansion of the streamlines. Lastly, cell-cell interactions lead to a slow recovery of the RBC distribution to its pre-compression distribution via cross-streamline migration if given sufficient length. We model the flow expansion of the RBC distribution with a logistic term and the cross-streamline recovery with an exponential decay:

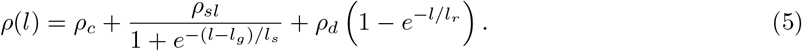

For *l* = 0, at the downstream end of the compression, the equation returns the ellipse radius inside the compression, which is within 5% of *ρ*_*c*_, due to the second term becoming very small when *l* = 0, but not vanishing. *ρ*_*sl*_ is the change in RBC distribution due to the flow expansion which occurs over a length scale 2*l*_*g*_. The length *l*_*s*_ determines the steepness of the slope. *ρ*_*d*_ is the change in RBC distribution due to cell-cell interaction, which occurs over a longer length scale *l*_*r*_ » *l*_*g*_. For long distances, *l* → ∞, we have *ρ*(*l*) = *ρ*_*c*_ + *ρ*_*sl*_ + *ρ*_*d*_ which is the width of the fully recovered RBC distribution and the width of the unperturbed RBC distribution before the compression. If *l* is not sufficiently large, the RBC distribution is still affected by the compression, and the RBC split in the bifurcation tends to be biased accordingly. Although it may be possible to construct a reduced-order model with fewer parameters, the advantage of Eq. (5) is that all parameters have a physical meaning and can potentially be predicted by separate models.

We obtained the numerical values of the parameters in the reduced-order model by fitting *ρ*(*l*) to the simulation data for 10% and 20% haematocrit, respectively. The parameters are listed in Table 2, and Figure 5d shows an excellent agreement between *ρ*(*l*) and the simulation data at *H*_*d*_ = 20%. In particular, we find that *l*_*g*_ ≈ 0.66*D*, independently of the chosen haematocrit. This value corresponds to the characteristic length which describes the streamline recovery after a distortion. We find that the cross-streamline recovery length *l*_*r*_ reduces by a factor of about 2 when the haematocrit is increased from 10% to 20%. This is in line with literature reporting that shear-induced diffusion is directly proportional to the particle concentration [46].

**Table 2.** Parameter values for Eq. (5) obtained by fitting the reduced-order model to simulation data.

| Haematocrit | $\rho_c$ | $\rho_{sl}$ | $\rho_d$ | $l_s$ | $l_g$ | $l_r$ |
| --- | --- | --- | --- | --- | --- | --- |
| 10% | 1.62 $\mu\text{m}$ | 6.40 $\mu\text{m}$ | 2.19 $\mu\text{m}$ | 0.17$D$ | 0.69$D$ | 13.8$D$ |
| 20% | 2.75 $\mu\text{m}$ | 7.41 $\mu\text{m}$ | 1.79 $\mu\text{m}$ | 0.16$D$ | 0.67$D$ | 5.1$D$ |

With the reduced-order model being calibrated, we can now predict the RBC partitioning at the downstream bifurcation and compare these results with actual RBC simulation data. We apply the separatrix model to the cross-sectional RBC distribution predicted by the reduced-order model at the length *l* that marks the distance between the compression and the point of bifurcation.

Table 3 compares the absolute difference in discharge haematocrit obtained from the HemeLB simulations and the reduced-order model. We find that the reduced-order model accurately predicts the impact of the compression on the RBC partitioning at the downstream bifurcation within 1% on average. Notwithstanding the assumptions underlying the reduced-order model, the relative error is low. Our novel approach, therefore, provides a means of modelling the disturbance caused by a compressed vessel in network simulations, which has not been possible using established empirical models [39, 41]. Despite this success, further simulations are necessary to extend the validity of the reduced-order model to a larger parameter space.

**Table 3.** Absolute differences of the discharge haematocrit in both child branches between the results of the HemeLB simulations and the predictions of the reduced-order model. In each box, the top left is the difference for the higher flowing child branch, whereas the bottom right is the difference for the lower flowing child branch. See Figure 1a–d for the respective geometries.

| Haematocrit | Control | Long compression<br>no recovery | Short compression<br>short recovery | Short compression<br>long recovery |
| --- | --- | --- | --- | --- |
| 10% | -0.02%<br>2.02% | 0.22%<br>0.42% | 0.00%<br>0.84% | 0.29%<br>-0.25% |
| 20% | 0.89%<br>-2.70% | -0.62%<br>3.23% | 0.29%<br>-0.61% | 0.96%<br>-2.54% |

### At 10% haematocrit, a converged suspension of RBCs requires a long development length

We observed that, at 20% haematocrit, the RMSD of the RBCs after 25*D* downstream of the compression has recovered to 98% of its original value prior to the compression (Figure 6a). However, at 10% haematocrit in the same geometry, the RMSD recovery is incomplete after the 25*D*. In fact, the reduced-order model predicts a partial recovery of the RMSD at 50*D* to only 91% of its pre-compression value (Figure 6b).

**Figure 6.**
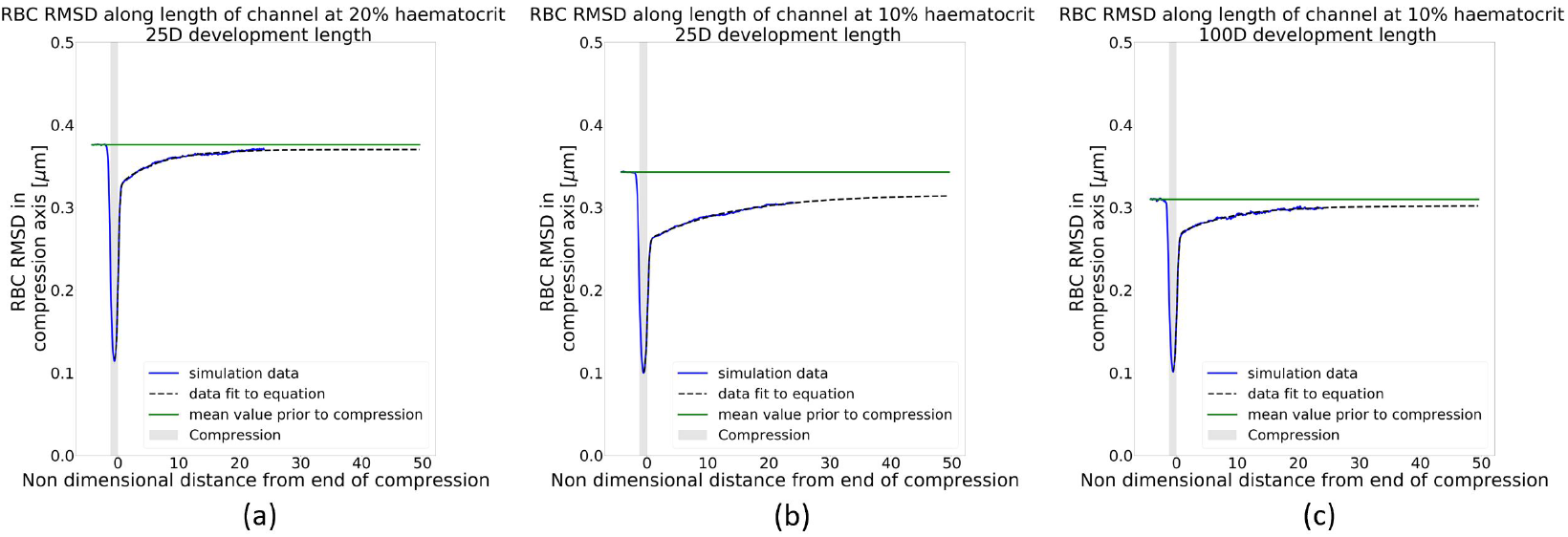
Recovery of RBC distribution after short compression. (a) Simulation at *H*_*d*_ = 20%, (b–c) simulations at *H*_*d*_ = 10%. (a) and (b) are simulations with the cells inserted after 25*D* of initialisation length. (c) is a simulation with cells inserted after an initialisation length of 100*D*. The blue line is the simulation data, the black line is the prediction of the reduced-order model, and the green line is the mean value of the RMSD prior to the compression.

Given the results in Figure 6b, we hypothesise that, at 10% haematocrit, details of the RBC initialisation in the simulation play a role. An inconsistent RBC distribution upstream of the compression might affect the overall outcome of the simulation. To confirm this, we increased the length of the periodic tube that is used to generate the RBC distribution fed into the compression geometry from 25*D* (Figure 6b) to 100*D* (Figure 6c). We observed that the longer tube leads to a narrowing in the pre-compression distribution of the RBCs (Figure 6c). At 10% haematocrit, when the initialisation length is 100*D*, we can assume that the RBC distribution has reached a steady state. In fact, Figure 6c shows that after 50*D* the RMSD of the RBCs has recovered to 98% of its pre-compression value. Despite the sensitivity of the pre-compression distribution on the cell initialisation strategy at *H*_*d*_ = 10%, we found that the RBC dynamics after the compression is quantitatively and qualitatively similar for both RBC initialisation lengths used.

## Discussion

The tumour microvasculature is abnormal and linked to tumour tissue hypoxia [1], which is a known biomarker for poor prognosis [2] and a barrier to recent promising immunotherapeutic approaches [3]. One such abnormality is vessel compression [5]. Previous studies have shown that decompressing tumour vessels leads to increased survival rates [8] via increased perfusion [1, 8] and oxygen homogenisation [9]. However, the mechanism linking tumour decompression to increased oxygen homogeneity is unclear. Oxygen binds to haemoglobin in red blood cells (RBCs) and is transported through the vasculature with the RBCs. We recently identified the reduced inter-bifurcation distance associated with the pro-angiogenic tumour environment as a source of oxygen heterogeneity via its impact on RBC splitting at bifurcations [14]. However, the impact that other tumour vascular phenotypes, such as vessel compression, play on this process is not known.

Motivated by the limitations on experimental methods available to query this process, we propose a computational model to elucidate the link between vessel compression and abnormal RBC partitioning at bifurcations. Our numerical simulations show that a vessel compression enhances the disproportional partitioning of RBCs at a downstream bifurcation in favour of the higher flow rate child branch. This is a consequence of the previously identified narrowing of the RBC distribution within the vessel cross section [17].

Similarly to previous studies [15, 17], we identify the mechanism leading to this narrowing as RBC cross-streamline migration towards the vessel centre due to an increased shear rate within the compression. Once the RBCs leave the compression, their cross-sectional distribution gradually goes back to their pre-compression configuration in a haematocrit-dependent manner. This process is significantly slower at 10% haematocrit, where the dynamics occur over a length of ∼ 50 vessel diameters. However, at 20% haematocrit, there is an almost instantaneous recovery, and after 2 vessel diameters in length, the difference in RBC partitioning compared to a control simulation is negligible. Furthermore, we show that at 30% haematocrit, the difference between the compressed and control simulation is negligible. This suggests the presence of a critical haematocrit above which vessel compression no longer alters the partitioning of RBCs at a bifurcation. We hypothesise that the different dynamics at 10%, 20% and 30% haematocrit are caused by cell-cell interaction which increases with haematocrit. We also show that an asymmetric compression does not lead to a measurable difference in the partitioning of RBCs compared to a symmetric compression. Likewise, a reduction and increase in flow rate by a factor of 5, respectively, does not significantly change the RBC partitioning.

We propose a reduced-order model to approximate the partitioning of RBCs at a bifurcation downstream of a compression, which we show has an error of 1% compared to our fully resolved numerical simulations, in the range of parameters studied. This model has the potential to overcome the computational tractability limitations associated with simulating RBC flow in large computational domains and will contribute to unravelling the dynamics of oxygen transport in large vascular tumour networks.

The implications of our findings are multiple. First, our results show that the effect of vessel compression on the downstream partitioning of RBCs is only apparent when discharge haematocrit and the distance between the compression and the bifurcation is sufficiently low. Another study by our group showed that two consecutive bifurcations within a short distance can also alter the partitioning of RBCs at the downstream bifurcation [14]. Furthermore, we showed that interbifurcation distances are much reduced in the tumour micro-environment, and Kamoun *et al*. showed that haemodiluted vessels are more common and are present across a larger range of diameters in tumour networks than in controls [12]. Taken together, this suggests that healthy vascular networks are structurally adapted to protect themselves from mechanisms leading to RBC transport heterogeneity and that this may be compromised in diseased networks.

Second, previous studies have shown that decompressing tumour vessels leads to increased survival rates [8, 9]. This has been attributed to a) increased tumour perfusion due to reduced vessel resistance [1, 8] and b) reduced hypoxia fraction and increased oxygen homogeneity [9]. Our results of anomalous RBC partitioning being unaffected by increases in flow rate support the view that increasing total perfusion through the network may not be sufficient to homogenise oxygenation if it is not accompanied by vessel decompression (or other forms of structural remodelling normalising RBC partitioning).

Third, Kamoun *et al*. found that up to 29% of tumour vessels in an animal model of glioma experience haemodilution (defined as having haematocrits below 5%) and proposed a mechanism whereby extravasated plasma from leaky vessels would be reabsorbed by other vessels and lead to haemodilution [12]. Along similar lines, recent studies have reported findings of tissue hypoxia near perfused vessels [2]. Our results demonstrate that, in the presence of vessel compression and uneven flow split at bifurcations, haematocrit can decrease from 20% to nearly 0% following two consecutive bifurcations without contributions from interstitial fluid. Our present findings, therefore, provide an alternative explanation of the occurrence of haemodilution in tumour networks. Future work should elucidate the relative importance of these two mechanisms.

Lastly, we identified that, in the semi-dilute regime of 10% haematocrit, achieving convergence in an RBC suspension that has been disturbed requires longer distances than previously thought. Katanov *et al*. reported that it takes a length of 25 diameters for the CFL of a randomly initialised suspension in a straight channel to converge [34]. However, our data suggest that up to 100*D* of length is required for an initially compacted RBC distribution to expand and reach a steady distribution when the haematocrit is 10%. This finding supports the view that the cross-sectional distribution of RBCs at low *in vivo* haematocrits may be away from equilibrium not only in diseased vascular networks, as we previously showed in tumours [14], but also under physiological conditions where inter-bifurcation lengths average fewer than 100*D*. Further research into the network-level dynamics arising from our results is warranted.

As a final comment, our results should be considered for *in vitro* experiments that need to carefully consider the design of microfluidic devices if full convergence of RBC suspensions is required in the semi-dilute regime. It is also an additional challenge for *in silico* studies with open boundary conditions, where not only the insertion of cells needs to be considered [47], but also their cross-sectional distribution.

## Conclusions

In this work we have demonstrated that vessel compression can alter RBC partitioning at a downstream bifurcation. Interestingly, this happens in a haematocrit-dependent and flow rate-independent manner. We argue that these findings contribute to the mechanistic understanding of haemodilution in tumour vascular networks and oxygen homogenisation following pharmacological solid tumour decompression. Furthermore, we have formulated a reduced-order model that will help future research elucidate how these effects propagate at a whole network level. Unravelling the causal relationship between tumour vascular structure and tissue oxygenation will pave the way for the development of new therapeutic strategies.

## Acknowledgments

We acknowledge the contributions of the HemeLB development team. Software development was supported by the Engineering and Physical Sciences Research Council (EPSRC) (grant eCSE-001-010). Supercomputing time on the ARCHER UK National Supercomputing Service (http://www.archer.ac.uk) was provided by the “UK Consortium on Mesoscale Engineering Sciences (UKCOMES)” under the EPSRC Grant No. EP/R029598/1. R.E. is funded by The University of Edinburgh through a Chancellor’s Fellow PhD studentship. D.H. is supported by the European Union’s Horizon 2020 research and innovation programme under grant agreement No 801423. M.O.B is supported by grants from EPSRC (EP/R021600/1, EP/R029598/1), Fondation Leducq (17 CVD 03), and the European Union’s Horizon 2020 research and innovation programme under grant agreement No 801423. T.K. and M.O.B. are supported by a grant from EPSRC (EP/T008806/1).

## Supplementary figures

**Figure S1.**
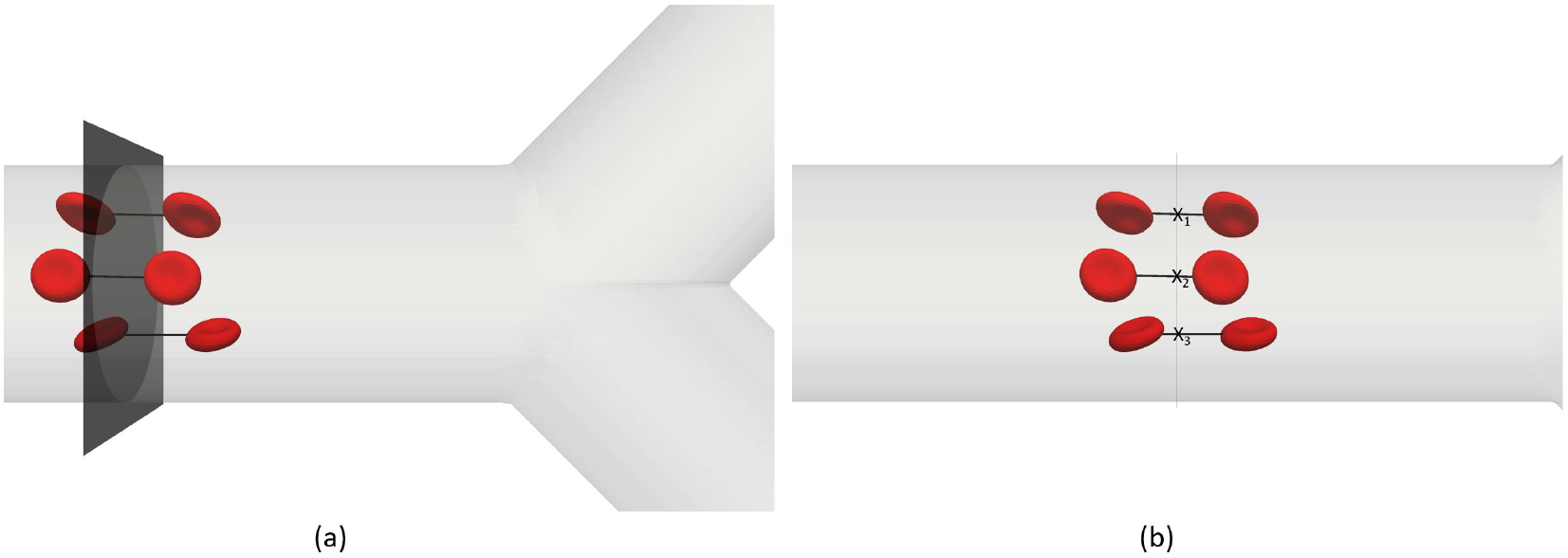
(a) Three RBCs before and after a plane of interest. Lines indicate RBC trajectories, assumed as straight. (b) Side view as each cell crosses the plane at a given coordinate (*x, y, z*). The RMSD is calculated in the compression axis (here seen as height of channel) by setting *x*_0_ as the channel centreline (always zero) and *x*_*i*_ as the height coordinate of the RBC as it crosses the plane. This measures the distribution in the height of the channel. For illustration purposes only three cells are shown, whereas several hundred are accounted for.

**Figure S2.**
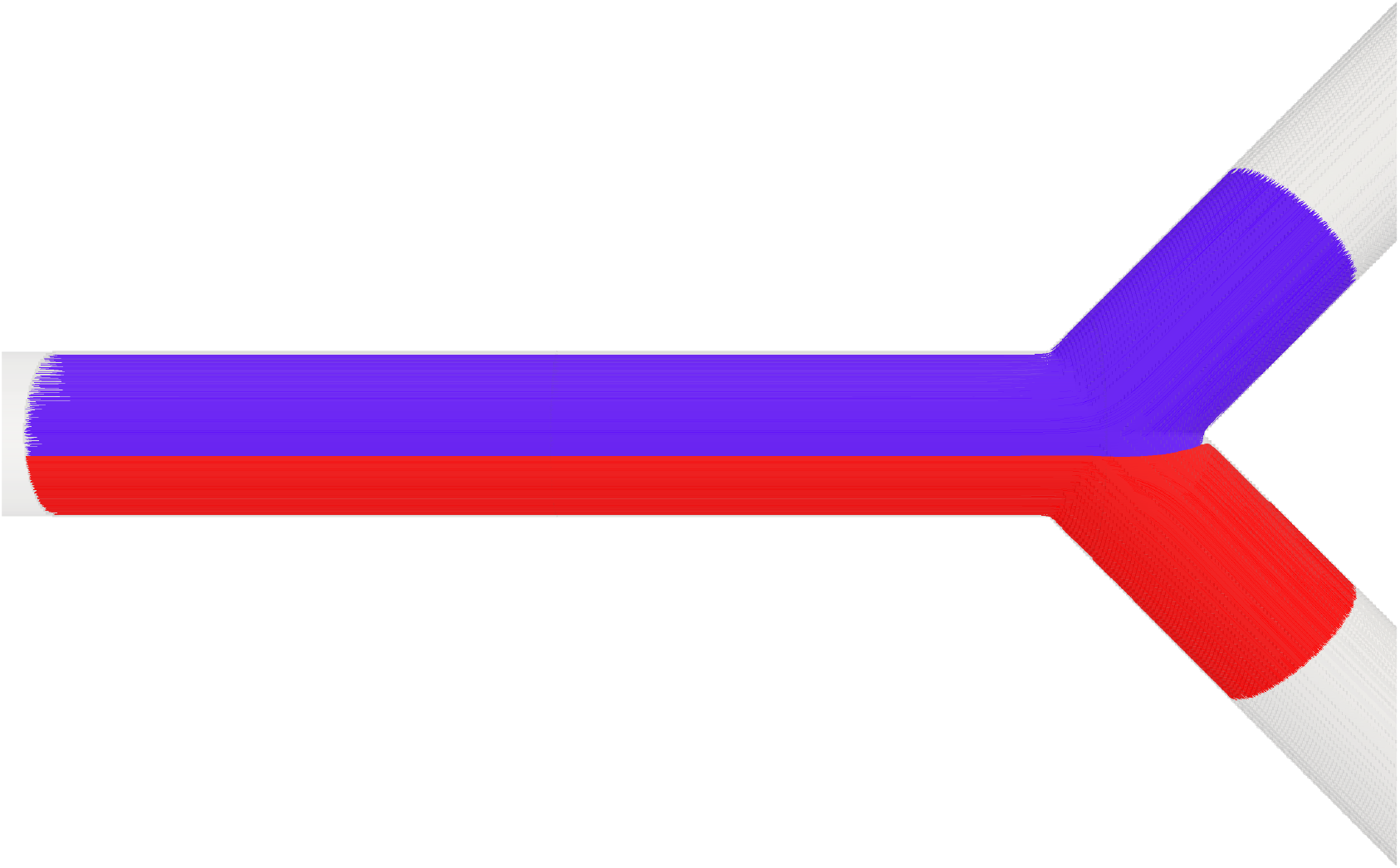
Intuition for separatrix. Blue/red lines are streamlines ending in the top/bottom child branch, respectively. The separatrix is the surface separating the blue from the red streamlines on the plane.

**Figure S3.**
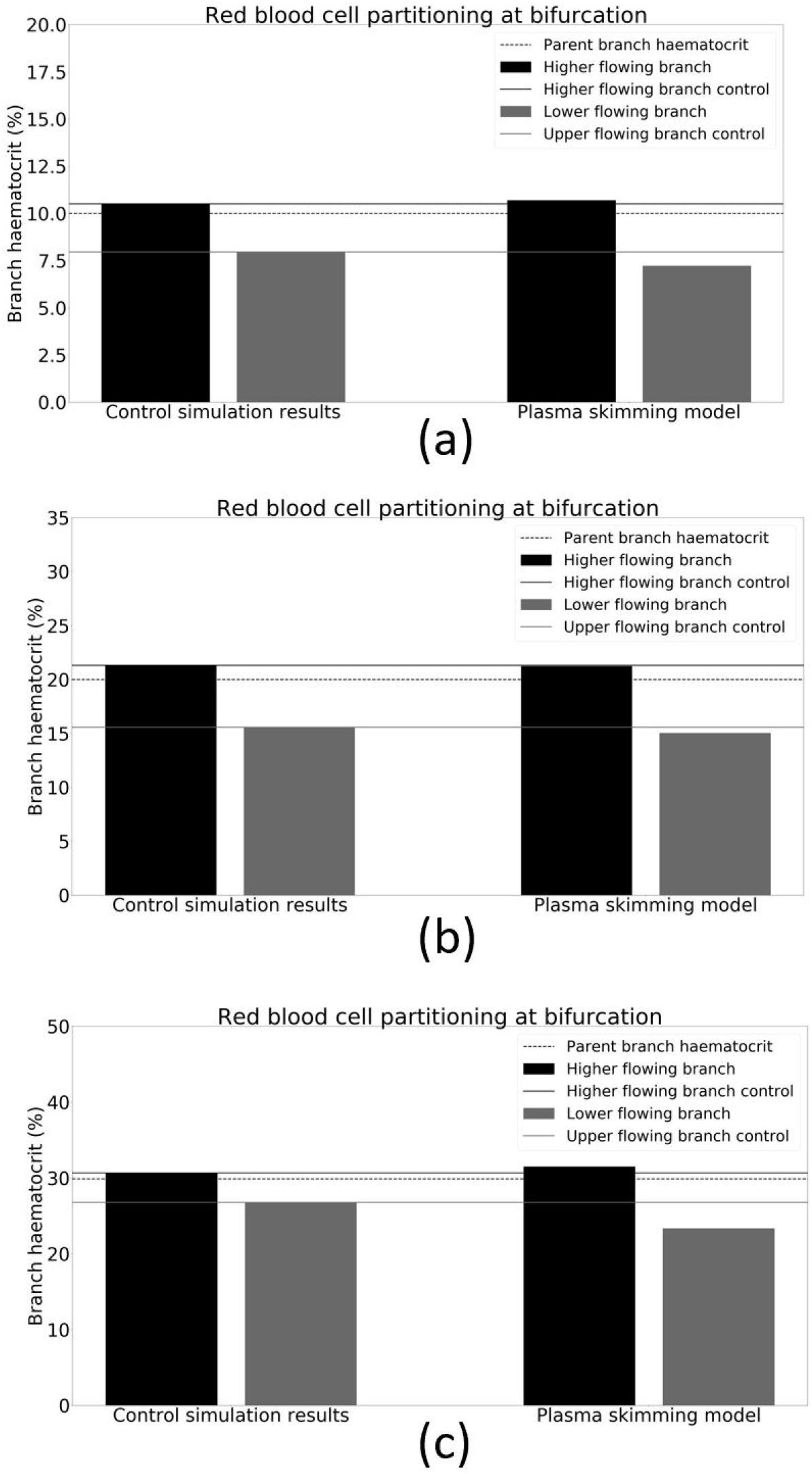
Comparison of simulation control data with empirical plasma skimming model [39, 40] with a flow ratio of 4. (a) Simulation at *H*_*d*_ = 10%. (b) Simulation at *H*_*d*_ = 20%. (c) Simulation at *H*_*d*_ = 30%.

**Figure S4.**
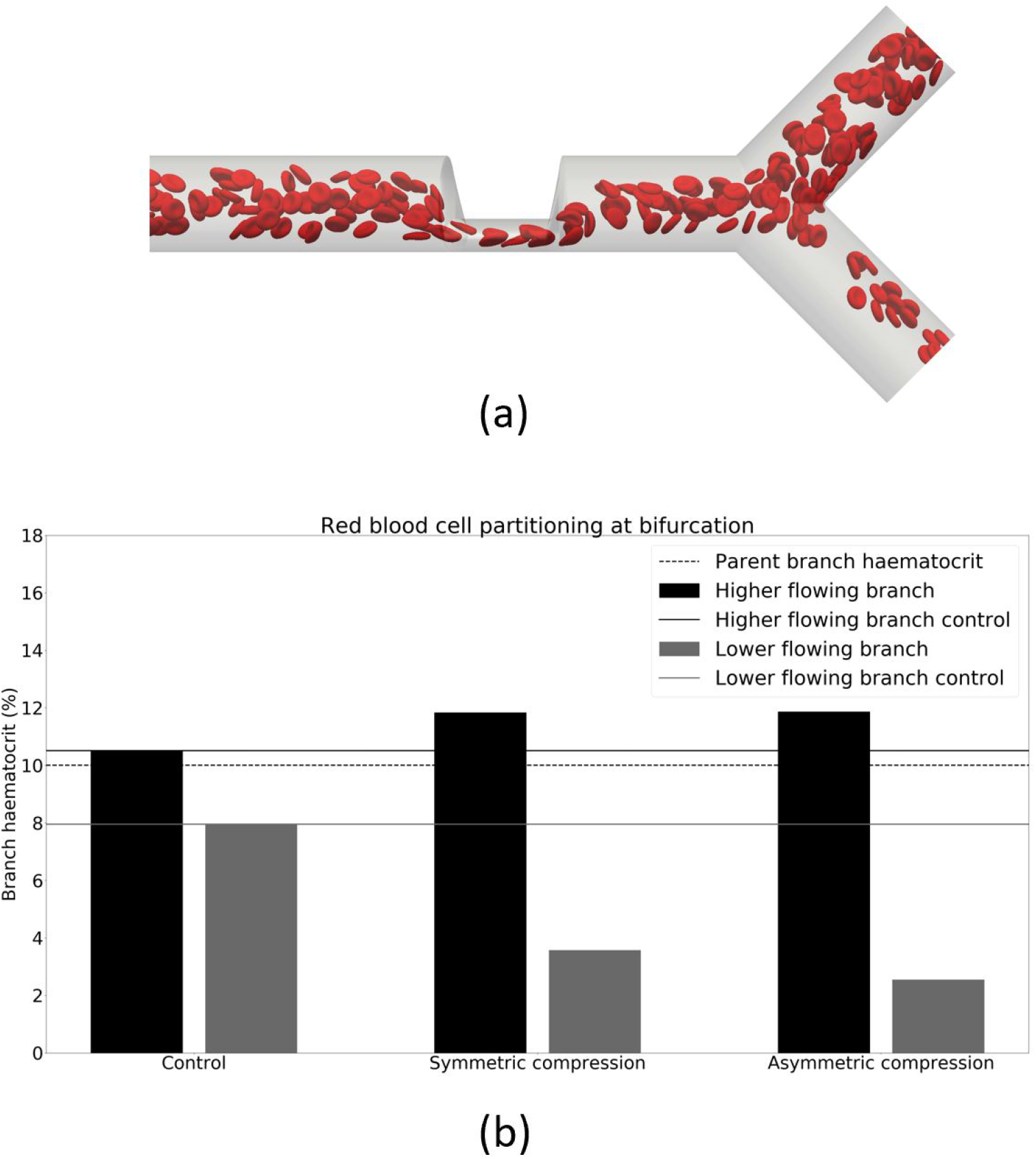
Phase separation of child branches after bifurcation at *H*_*d*_ = 10% comparing effect of compression asymmetry. (a) Snapshot of the asymmetric short geometry with the same dimensions as the short geometry. (b) From left to right are the haematocrit of the child branches for a control geometry, a symmetric compression, and an asymmetric compression (a). Results show a negligible difference between a symmetric and asymmetric geometry. In black is the higher flowing child branch and in grey the lower flowing child branch. The solid lines are the control discharge haematocrits. The dotted line illustrates the discharge haematocrit of the parent branch.

**Figure S5.**
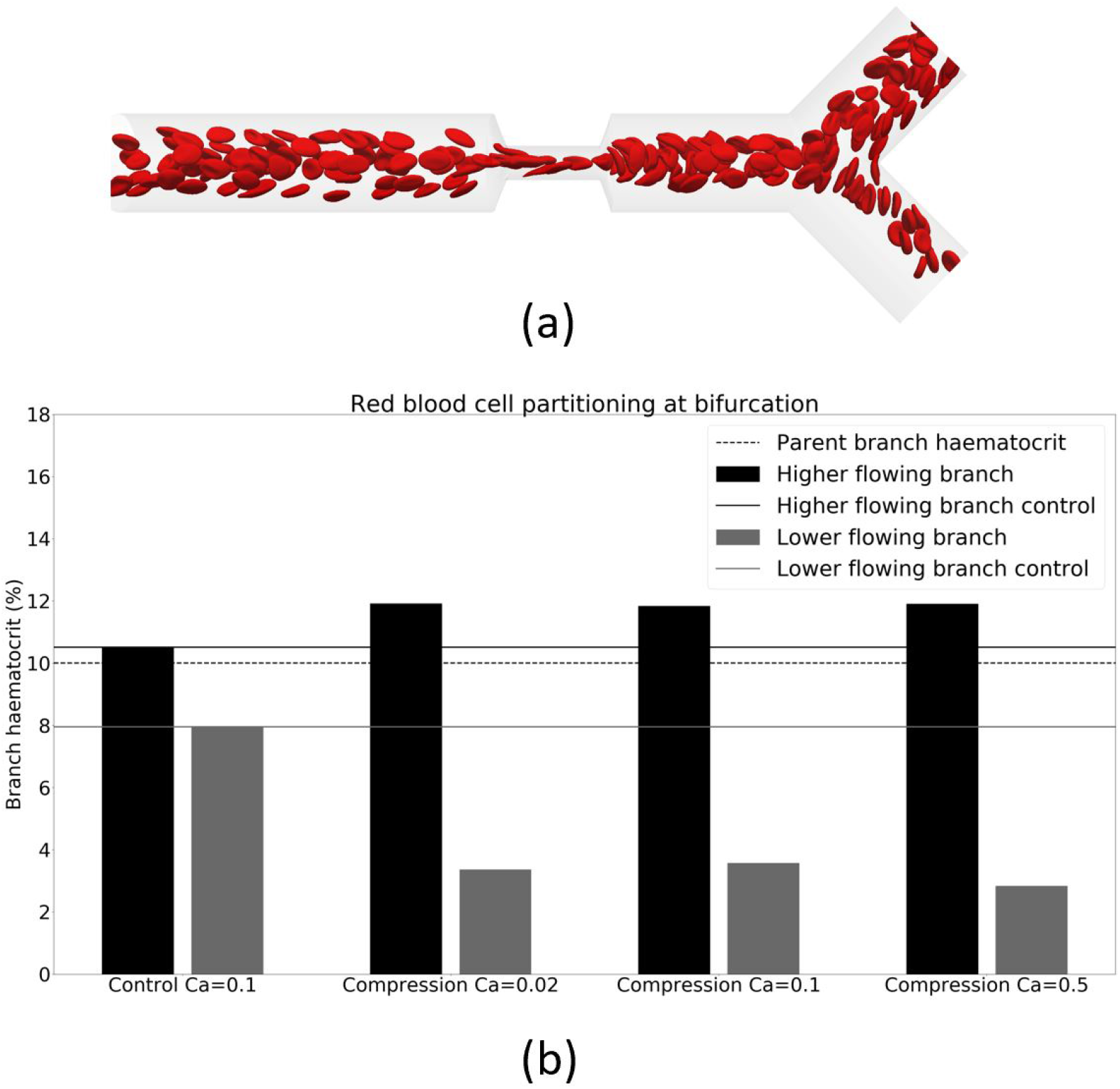
Phase separation of child branches after bifurcation at *H*_*d*_ = 10%, comparing effect of different channel flow rates (increasing capillary number denotes increasing flow rate). (a) Snapshot of the short compression with a higher channel flow rate and a capillary number of 0.5. (b) Haematocrit of the child branches for a control geometry, on the left, and a compression geometry (a) at three different capillary numbers. In black is the higher flowing child branch and in grey the lower flowing child branch. The solid lines are the control discharge haematocrits. The dotted line illustrates the discharge haematocrit of the parent branch.

